# TDbasedUFE and TDbasedUFEadv: bioconductor packages to perform tensor decomposition based unsupervised feature extraction

**DOI:** 10.1101/2023.05.14.540687

**Authors:** Y-h. Taguchi, Turki Turki

## Abstract

**Motivation:** Tensor decomposition (TD) based unsupervised feature extraction (FE) was proposed almost five years ago and has been successfully applied to a wide range of bioinformatics problems ranging from biomarker identification to the identification of disease-causing genes and drug repositioning. Despite its successful applications, the use of TD-based unsupervised FE was not widely accepted because of the unpopularity of TD in this field.

**Results:** To overcome this difficulty, we developed two bioconductor packages, TDbasedUFE and TDbasedUFEadv. Using these two packages, all researchers who are not familiar with the concept of TD can make use of TD-based unsupervised FE for their purposes. When the performances of two specific functions, identification of differentially expressed genes and multiomics analysis, are implemented in TDbasedUFE and compared with those of two state-of-the-art (SOTA) methods (i.e., DESeq2 and DIABLO), TDbasedUFE can outperform these two SOTAs.

**Availability and implementation:** TDbasedUFE and TDbasedUFEadv are freely available as R/Bioconductor packages hosted at https://bioconductor.org/packages/TDbasedUFE and https://bioconductor.org/packages/TDbasedUFEadv, respectively.

## Introduction

Tensor decomposition (TD) based unsupervised feature extraction (FE) was proposed several years ago [Taguchi, 2017] and was applied to a wide range of problems [Taguchi, 2020]. Despite various successful applications, TD-based unsupervised FE has never been widely employed, possibly because of the unpopularity of TD in this field. We have developed two bioconductor packages, TDbasedUFE and TDbasedUFEadv, so researchers can use TD-based unsupervised FE easily even without detailed knowledge of TD. In this application note, we introduce these two packages to researchers.

### Features

TD-based unsupervised FE [Taguchi, 2017] was developed from principal component analysis (PCA) based unsupervised FE [Taguchi and Murakami, 2013] that was proposed ten years ago. As the complexity of datasets increases as multiple measurement conditions came to be applied to individual studies (e.g., comparisons of multiple tissues from human subjects instead of those from human patients restricted to a single tissue), tensors which can have multiple indices, each of which can have multiple comparison criteria, were employed instead of matrices. For example, three mode tensor *x*_*ijk*_ can naturally store the expression of *i*th gene at *k*th tissue of *j*th human subjects. If we are forced to use a matrix that has only two indices corresponding to rows and columns instead of tensors, the tissue index and the human index must be mixed up into a column, which is harder to interpret than tensors.

TDbasedUFE and TDbasedUFEadv are easy for a person who is not familiar with the concept of tensors to use. Since the matrix can be regarded as a two-mode tensor, TDbasedUFE and TDbasedUFEadv are also used to apply PCA-based unsupervised FE to the dataset. TDbasedUFE focuses only on two popular functions among those possible by TD-based unsupervised FE, since TD-based unsupervised FE can perform numerous applications, not all of which are required by the majority of people. Two popular functions implemented in TDbasedUFE are the identification of differentially expressed genes (DEGs) and multiomics analyses. For the DEG identification, a basic algorithm is based upon a recent paper [Taguchi and Turki, 2022a] where a newly implemented standard deviation (SD) optimization was established. For multiomics analysis, although a basic algorithm is based upon the paper [Taguchi and Turki, 2022c], SD optimization, which was not invented at the time the study was published, was considered in the implementation of multiomics analysis in TDbasedUFE. Although it is not specifically implemented for application to DNA methylation profiles, we found that the algorithm found in the paper [Taguchi and Turki, 2022a] is also applicable to DNA methylation profiles as it is [Taguchi and Turki, 2023]. In this sense, any other differential analysis of single omics is supposed to be performed by functions implemented in TDbasedUFE. In actuality, we have found [Roy and Taguchi, 2022] that histone modification profiles can be treated by the algorithm found in the paper [Taguchi and Turki, 2022a].

TDbasedUFE and TDbasedUFEadv accept a multiple omics profile dataset formatted as a tensor to which TD is applied (we employed Tucker decomposition as a TD using the higher-order singular value decomposition (HOSVD) [Taguchi, 2020] algorithm). For example, if *x*_*ijk*_ ∈ ℝ ^*N* × *M* × *K*^ represents the gene expression of *i*th gene of *j*th human subject’s *k*th tissue (Fig. 1 left), TD is applied to *x*_*i j k*_ and we get

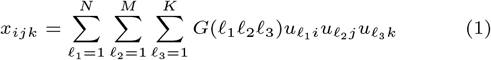

where *G* ∈ ℝ^*N* × *M* × *K*^ is a core tensor that represents the weight of the product 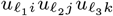 to *x*_*ijk*_ and 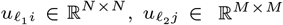, and 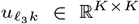 are singular value matrices and orthogonal matrices. At first, singular value vector attribute to samples 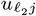 and 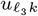 are investigated to identify those of interest (e.g.,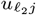 represents the distinction between healthy controls and patients and 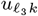 represents tissue specificity (e.g., expressive only in the heart)). Then, singular value vectors attributed to genes (i.e., features) 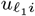 that share *G* of the largest absolute value with the identified 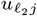 and 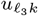 are selected. Features (*i*s) with larger absolute values of 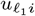 are identified based upon *P* -values computed by assuming that 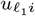 obeys Gaussian distribution (null hypothesis) as

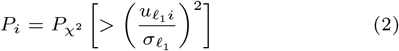

where 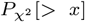 is the cumulative *χ*^2^ distribution where the argument is larger than *x* and 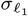 is the optimized standard deviation such that 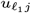 obeys Gaussian distribution as much as possible (see [Taguchi and Turki, 2022a] for more details about how to optimize 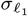). Then *P*_*i*_s are adjusted by Benjamini-Hochberg criterion to consider multiple comparison correction. Finally, is with adjusted *P*_*i*_ less than threshold value (typically, 0.01) are selected.

**Fig. 1.**
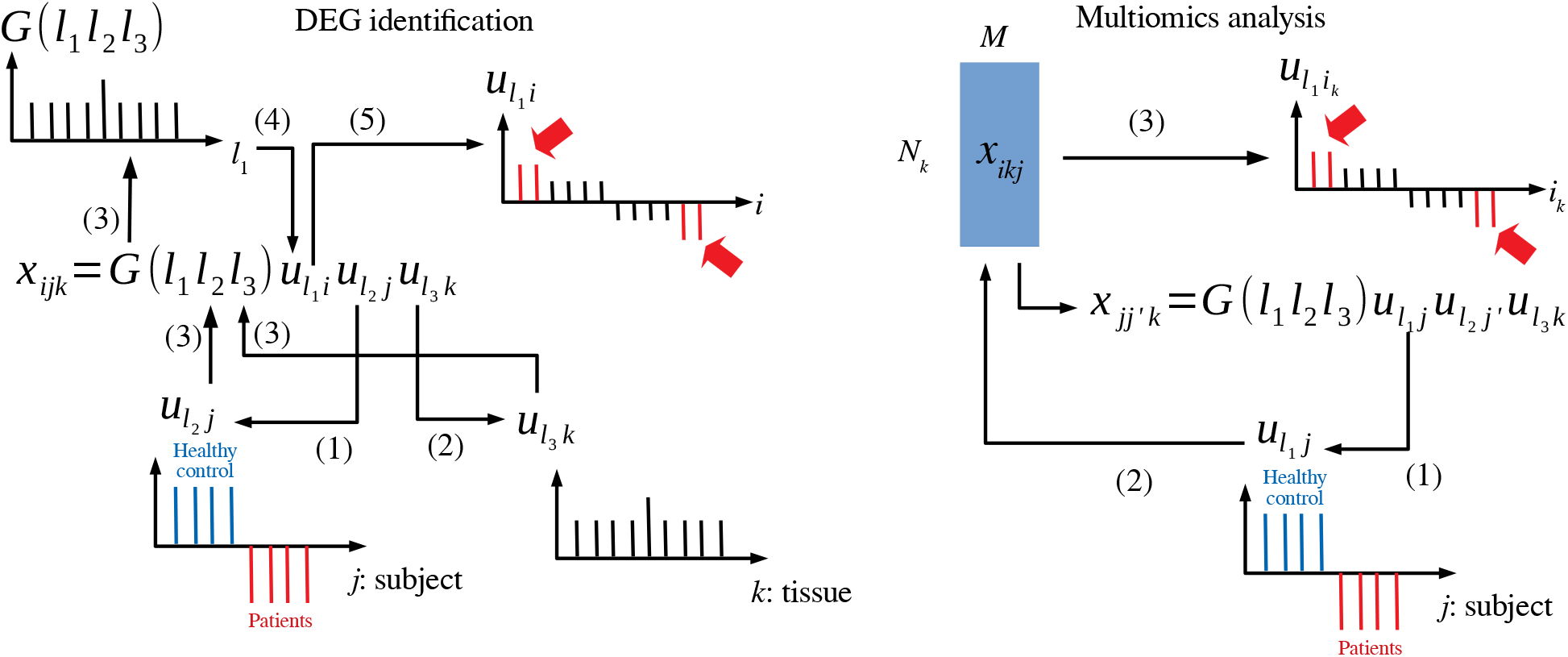
Schematic figure that explains TD-based unsupervised FE. Left: DEG identification, (1)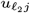 associated with the distinction between patients and healthy controls is selected. (2) 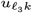 associated with tissue specificity is selected. (3) Investigate *G*(*ℓ*_1_*ℓ*_2_*ℓ*_3_) with fixed *ℓ*_2_ and *ℓ*_3_. (4) 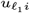 with *G* of the largest absolute value is selected. (5) *i*s (indicated in red) whose absolute values are significantly larger than expected are selected. Right: Multiomics analysis, (1) 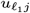 associated with the distinction between patients and healthy controls is selected. (2)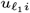 is computed from 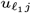 (3) *i*s (indicated in red) whose absolute values are significantly larger than expected are selected.

On the other hand, when TDbasedUFE was applied to multiomics datasets (Fig. 1 right), the multiomics profiles are formatted as 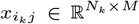 (i.e., *k*th omics datasets are associated with as many as *N*_*k*_ features). 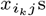 are multiplied with each other and got

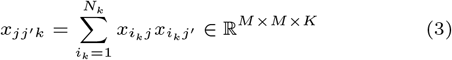

to which HOSVD was applied as

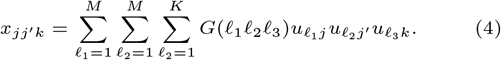

After identifying 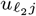coincident with labels (e.g., patients and healthy control), singular value vectors attributed to individual features associated with *k*th omics are computed as

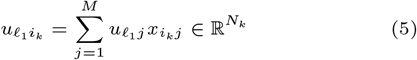

and 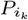 is computed as

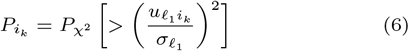

and *i*_*k*_ s associated with adjusted 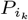 less than 0.01 are selected.

In contrast to TDbasedUFE that can perform only two tasks, TDbasedUFEadv can perform more complicated tasks. For example, TDbasedUFEadv can perform [Ng and Taguchi, 2020] integrated analysis of two omics profiles that share samples differently from TDbasedUFE and reduce the memory required by summing up the samples index. TDbasedUFEadv also can perform integrated analysis of two omics profiles that share features [Taguchi and Turki, 2019]. TDbasedUFEadv can also perform integrated analysis of multiple (more than two) omics profiles sharing either feature [Taguchi and Turki, 2022b] or samples [Taguchi and Turki, 2021] differently from TDbasedUFE.

## Results

See the supplementary materials for the full list of identified features as well as the results of the enrichment analysis in this section. For more details, see the supplementary document.

Although there are many applications proposed since the publication of our book [Taguchi, 2020], a few examples are shown here to demonstrate the usefulness of TDbasedUFE. We consider the ACC.rnaseq data from RTCGA.rneseq [Kosinski, 2023] package in Bioconductor. The labels used to select singular value vectors attributed to samples and coincident with labels are patient.stage event.pathologic stage composed of four classes (“stage i” to “stage iv”). A tensor *x*_*ijk*_ ∈ ℝ ^*N* × 9 × 4^ represents the expression of *i*th gene of *j*th replicates of *k*th stage. HOSVD was applied to *x*_*ijk*_ and we get TD as in eq. (1) (see the supplementary document for the R code to perform DEG identification using TDbasedUFE). Since 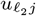 is attributed to replicates, 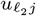 is expected to have constant value regardless of how *j* and *ℓ*_*2*_ = 1 turned out to satisfy this requirement (Fig. S1 left). On the other hand, 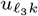 is expected to have monotonic dependence on *k* (Fig. S1 right); *ℓ*_3_ = 3 is most coincident with monotonic dependence upon *k*. Once *ℓ*_*2*_ and *ℓ*_*3*_ are selected by the user with the interactive interface, TDbasedUFE automatically selects 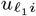 with which *i*s are selected. As a result, 1,692 genes are selected with the threshold adjusted *P* -values of 0.01.

To evaluate TDbasedUFE’s ability to select genes, we apply DESeq2 [Love et al., 2014], a state-of-the-art method to *x*_*ijk*_, although DESeq2 is not applied to *x*_*ijk*_ but to the unfolded matrix *x*_*i*(*jk*)_ ∈ ℝ ^*N* × 36^ where *j* and *k* are merged into a column index (see the supplementary document for the R code to perform DEG identification using DESeq2). Then we get as few as 138 genes associated with adjusted *P* -values less than 0.01. Thus, from the perspective of the number of identified DEGs, TDbasedUFE is clearly superior to DESeq2.

Although TDbasedUFE could identify more DEGs than DESeq2, if identified genes are not biologically reasonable, a higher number of identified DEGs is meaningless. To see if DEGs selected by TDbasedUFE is more biologically reasonable than those identified by DESeq2, we employed the enrichR [Jawaid, 2023] package in CRAN as demonstrated in the vignette entitled “Enrichment” in the TDbasedUFEadv package considering the “KEGG 2021 HUMAN,” “GO Molecular Function 2015,” “GO Cellular Component 2015,” and “GO Biological Process 2015” categories. Then we found that 129, 151, 143, and 923 terms are associated with adjusted *P* -values less than 0.05 when 1,692 genes selected by TDbasedUFE are considered. On the other hand, when 138 genes selected by DESeq2 are considered, 0, 0, 3, and 12 terms are associated with adjusted *P* -values less than 0.05. Thus, from the perspective of the number of biological terms identified, TDbasedUFE is superior to DESeq2.

To demonstrate TDbasedUFE’s ability applied to a multiomics dataset, we employ the curatedTCGA [Ramos et al., 2020] package to retrieve profiles other than the gene expression of the ACC dataset in TCGA (see the supplementary document for the R code to perform DEG identification using TDbasedUFE). We have collected miRNA 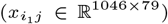, gene expression 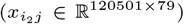, and methylation 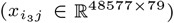 from curatedTCGA and applied TDbasedUFE to these three profiles as a mulitomics dataset. After applying HOSVD to the generated tensor *x*_*jj*′ *k*_ ∈ ℝ^79 × 79 × 3^, we found that 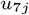 (Fig. S2 upper) is associated with the distinction between four stages and 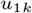 (Fig. S2 lower) is constant regardless to *k* (i.e., omics). 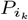 is attributed to *i*_*k*_ by eq. (6) using 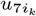 generated from 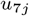 by eq. (5). After correcting 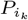, we have found 23 miRNAs among 1,046 miRNAs, 1,016 mRNAs among 20,501 mRNAs, and 7,295 probes among 485,577 methylation probes are associated with adjusted 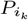 less than 0.01 (these features are expected to be distinct between the four stages as well).

To compare TDbasedUFE’s performance with other SOTA methods, we employed DIABLO implemented in the mixomics package [Rohart et al., 2017] in Bioconductor (see the supplementary document for the R code to perform mulitiomics analysis using DIABLO). Although we have employed the minimum setup (folds=2, nrepeat=1) since DIABLO did not finish even within three hours when the recommended setup in the vignette (folds=10, nrepeat=10) was employed, DIABLO did not converge to the solution with few enough errors up to ten components (ncomp=10) and showed no tendency that errors decrease as the number of components increases (Fig. S3). Thus, we gave up employing DIABLO applied to the current multiomics dataset since DIABLO did not allow us to select features if not converged to the solution with few enough errors; TDbasedUFE is obviously superior to a SOTA, DIABLO.

To evaluate these miRNAs, mRNAs, and methylation probes identified by TDbasedUFE biologically, we have uploaded those identified to various databases. At first, we uploaded the identified miRNAs to DIANA-mirpath v3.0 [Vlachos et al., 2015] and found many cancer-related KEGG pathways are enriched (see the supplementary document for URL to DIANA-mirpath using these miRNAs). Next, we uploaded the identified mRNAs to Enrichr [Xie et al., 2021], and found many cancer-related ones in the “KEGG 2021 Human” categories and various cancer cell lines. Finally, we uploaded 2,668 unique gene symbols associated with the identified 7,295 probes to Enrichr and found some cancer-related ones in “KEGG 2021 Human” and various cancer cell lines. We can conclude that the identified miRNAs, mRNAs, and methylation probes identified are biologically reasonable.

## Conclusions

In this application note, we have introduced TDbasedUFE and TDbasedUFEadv, which can perform TD-based unsupervised FE without much knowledge of tensor decompositions. They can outperform two SOTA, DESeq2 and DIABLO, when they are applied to DEGs identification and multiomics analysis, respectively. These two packages enable users to use TD-based unsupervised FE easily.

## Author contributions statement

Y. H. T. and T. T. wrote an original paper and reviewed the paper. Y. H. T. has developed the package and performed analysis. T. T. and Y. H. T. validated the results.

## Acknowledgments

This work is supported in part by funds from the Chuo University (TOKUTEI KADAI KENKYU).

