## Supplementary material for "TDbasedUFE and TDbasedUFEadv: bioconductor packages to perform tensor decomposition based unsupervised feature extraction": Suppl_doc.pdf

**3 Supplementary Figures** **3**

**4 Supplementary Data** **3**

### 1 DEG identification

#### 1.1 The R code to perform DEG identification by TDbasedUFE

```
library(RTCGA.rnaseq)
library(RTCGA.clinical)
library(TDbasedUFE)
stage <- sort(unique(ACC.clinical$patient.stage_event.pathologic_stage))
index1 <- ACC.clinical$patient.bcr_patient_barcode %in%
  tolower(substring(ACC.rnaseq[,1],1,12))
index0 <- NULL
for (i in seq_len(4)){
  index0 <- cbind(index0,
                 which(ACC.clinical$patient.stage_event.pathologic_stage[index1]
                      ==stage[i])[seq_len(9)])
}
index2<- matrix(match(ACC.clinical[index1,][index0,10],
      tolower(substring(ACC.rnaseq[,1],1,12))),ncol=4)
Z <- array(t(ACC.rnaseq[index2,-1]),c(dim(ACC.rnaseq)[2]-1,9,4))
Z <- PrepareSummarizedExperimentTensor(matrix(rep(stage,each=9),9,4),colnames(ACC.rnaseq)[-1],Z)
HOSVD <- computeHosvd(Z)
input_all <- selectSingularValueVectorSmall(HOSVD)
index <- selectFeature(HOSVD,input_all)
head(tableFeatures(Z,index))
table(tableFeatures(Z,index)$"adjusted p value"<0.01)
```

#### 1.2 The R code to perform DEG identification by DESeq2

```
library(DESeq2)
sampleTable <- data.frame(
  sampleName=ACC.clinical$patient.bcr_patient_barcode[index1][index0],
  fileName =rep("",36),
  condition=rep(stage,each=9))
sampleTablecondition <- data.frame(condition=factor(gsub(" ", "_",sampleTable$condition)))
names(sampleTablecondition) <- "condition"
X <- Z@value
```

```
dim(X) <- c(dim(X)[1],36)
rownames(X) <- colnames(ACC.rnaseq)[-1]
dds <- DESeqDataSetFromMatrix(countData =trunc(X),
                              colData = sampleTablecondition,
                              design= ~  condition)

dds <- DESeq(dds)
res <- results(dds)
res1 <- res[!is.na(res$padj),]
table(res1$padj<0.01)
```

### 2 Multiomics analysis

#### 2.1 The R code to perform multiomics analysis by TDbasedUFE

```
library(curatedTCGAData)
library(MultiAssayExperiment)
library(TCGAutils)
(accmae <- curatedTCGAData(
     "ACC", c("*RNASeq*", "*Methylation*"), version = "2.0.1", dry.run = FALSE
 ))
 PID1 <- substring(colnames(accmae[[1]]),1,12)
PID2 <- substring(colnames(accmae[[2]]),1,12)
PID4 <- substring(colnames(accmae[[4]]),1,12)
accmae[[1]] <- accmae[[1]][,match(PID2,PID1)]
accmae[[4]] <- accmae[[4]][,match(PID2,PID1)]
stage <- colData(accmae)$pathologic_stage[match(PID2,rownames(colData(accmae)))]
Z <- PrepareSummarizedExperimentTensorSquare(
  sample=matrix(PID2,ncol=1),
  feature=list(miRNA=rownames(accmae[[1]]),
             RNAseq=rownames(accmae[[2]]),
             Methylation=rownames(accmae[[4]])),
  value=convertSquare(list(data.matrix(accmae[[1]]@assays@data@listData[[1]]),
                           data.matrix(accmae[[2]]@assays@data@listData[[1]]),
                           data.matrix(accmae[[4]]@assays@data@listData[[1]]))),
  sampleData=list(matrix(stage,ncol=1)))
HOSVD <- computeHosvdSqure(Z)
cond <- list(attr(Z,"sampleData")[[1]],attr(Z,"sampleData")[[1]],seq_len(3))
input_all <- selectSingularValueVectorLarge(HOSVD,cond)
index <- selectFeatureSquare(HOSVD,input_all,list(data.matrix(accmae[[1]]@assays@data@listData[[1]]),
                                                   data.matrix(accmae[[2]]@assays@data@listData[[1]]),
                                                   data.matrix(accmae[[4]]@assays@data@listData[[1]])),
                              de=c(0.1,0.03,0.03),interact=FALSE)
for (id in seq_len(3)){print(head(tableFeaturesSquare(Z,index,id)))}
```

#### 2.2 URL for DIANA-mirpath ver. 3.0

Link to DIANA-mirpath ver 3.0 using identified miRNAs

#### 2.3 The R code to perform multiomics analysis by DIABLO

```
    X <- list(miRNA=t(data.matrix(accmae[[1]]@assays@data@listData[[1]])),
          mRNA=t(data.matrix(accmae[[2]]@assays@data@listData[[1]])),
          metyl=t(data.matrix(accmae[[4]]@assays@data@listData[[1]])))
dimnames(X[[3]])[[1]] <- dimnames(X[[1]])[[1]]
dimnames(X[[3]])[[2]] <- accmae[[4]]@NAMES
for(i in seq_len(3))
```

```
{
  rownames(X[[i]]) <- substring(rownames(X[[i]]),1,12)
}
Y <- stage
Y[is.na(Y)] <- "no stage"
design <- matrix(0.1, ncol = length(X), nrow = length(X),
                 dimnames = list(names(X), names(X)))
diag(design) <- 0
design
diablo.tcga <- block.plsda(X, Y, ncomp = 10, design = design,near.zero.var = TRUE)
set.seed(123) # For reproducibility, remove for your analyses
perf.diablo.tcga = perf(diablo.tcga, validation = 'Mfold', folds = 2)
plot(perf.diablo.tcga,ylim=c(0,5))
```

### 3 Supplementary Figures

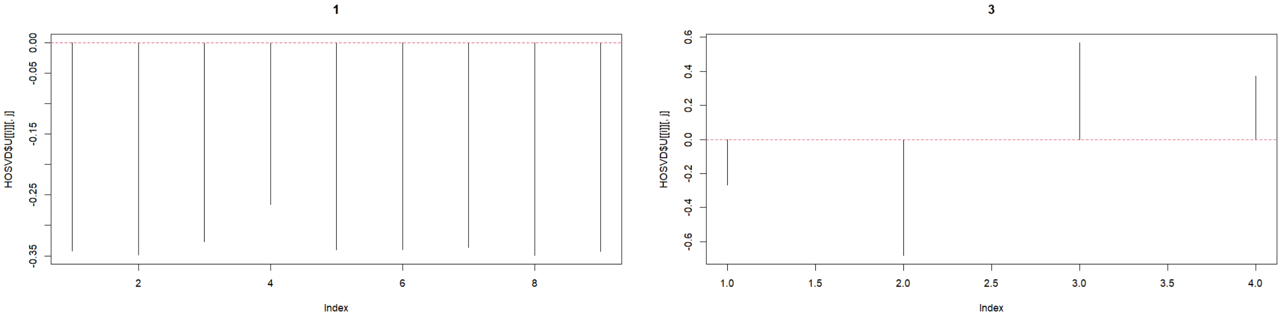

Figure S1: Left: $u_{1j}$, right: $u_{3k}$

### 4 Supplementary Data

- gene_DEG: Genes selected by TDbasedUFE for DEG identification and associated enrichment analysis (KEGG, GO BP, GO MF, GO CC)
- gene_DESeq2: Genes selected by DEseq2 for DEG identification and associated enrichment analysis (KEGG, GO BP, GO MF, GO CC)
- gene_multi: Genes selected by TDbasedUFE for multiomocs analysis and associated enrichment analysis (KEGG, GO BP, GO MF, GO CC, ARCHS4 Cell-lines, NCI-60 Cancer Cell Lines)
- methyl_multi: Genes associated with methylation probes selected by TDbasedUFE for multiomocs analysis and associated enrichment analysis (KEGG, GO BP, GO MF, GO CC, ARCHS4 Cell-lines, NCI-60 Cancer Cell Lines)
- miRNAs_multi: miRNAs selected by TDbasedUFE for multiomocs analysis

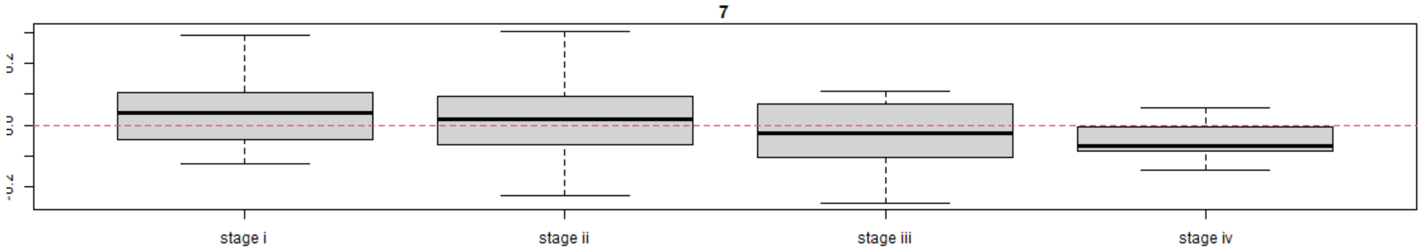

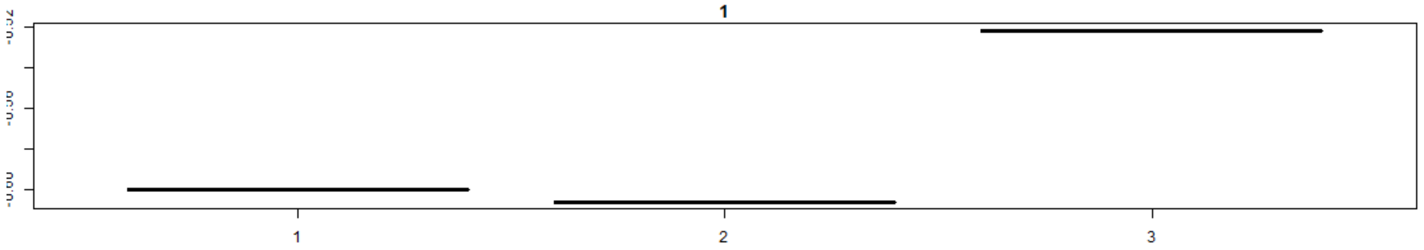

Figure S2: Upper: $u_{7j}$, lower: $u_{1k}$

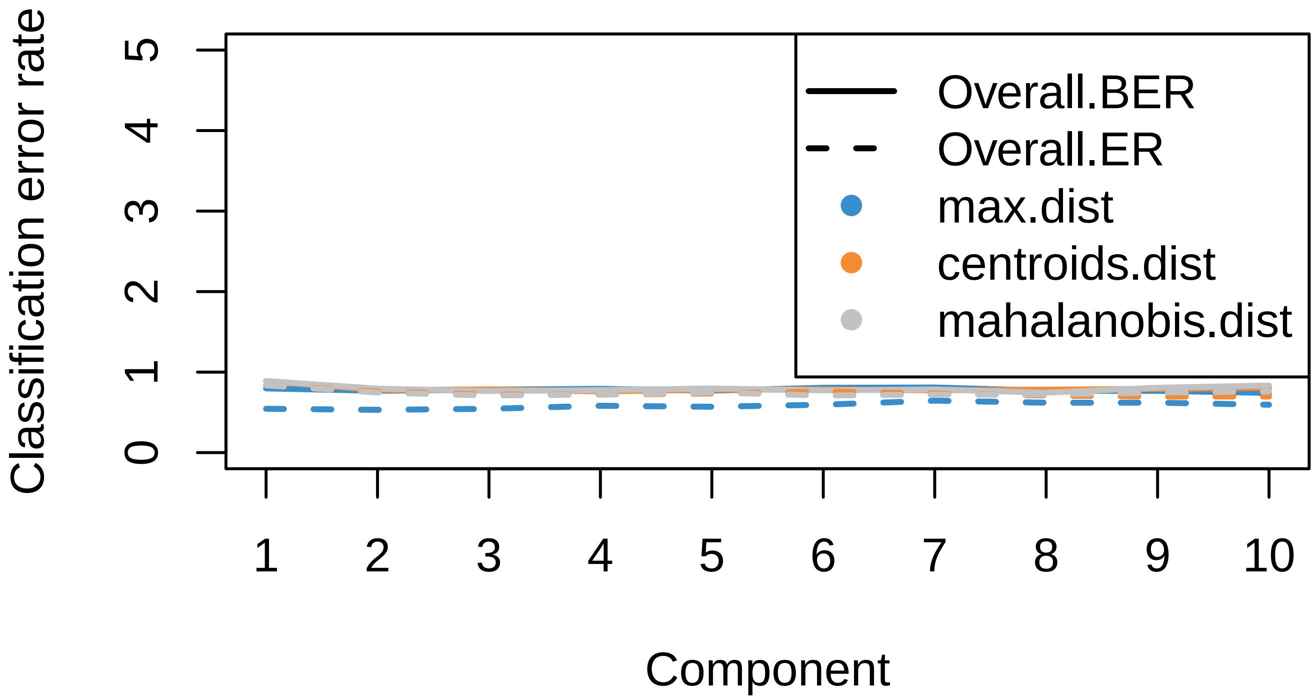

Figure S3: Dependence of errors upon the number of component for DIABLO. Please note that upper limit of vertical axis is taken to be 500 % errors to avoid overlap of legends with plot. Thus the amount of errors are as large as 50 %.
